# Identification of Polynucleotide Phosphorylase (PNPase) in *Escherichia coli* Involved in Persister Formation

**DOI:** 10.1101/310987

**Authors:** Nan Wu, Yumeng Zhang, Shanshan Zhang, Youhua Yuan, Shuang Liu, Peng Cui, Wenhong Zhang, Ying Zhang

## Abstract

Despite the identification of many genes and pathways involved in the persistence phenomenon of bacteria, the mechanisms of persistence are not well understood. Here, using *Escherichia coli* as a model, we identified polynucleotide phosphorylase (PNPase) as a key regulator in persister formation. We successfully constructed *pnp* knockout mutant strain and its complemented strain, and exposed the *pnp* knockout mutant and complemented strain to antibiotics and stress conditions. The results showed that, compared with the wild-type W3110, the *pnp* knockout strain had defect in persistence to antibiotics and stress conditions, and the persistence to antibiotics and stresses was restored upon complementation. RNA-Seq was performed to identify the transcriptome profile in the *pnp* knockout strain compared with wild-type strain W3110, and the data revealed that 242 (166 up-regulated, and 76 down-regulated) genes were differentially expressed in the *pnp* knockout mutant strain. KEGG pathway analysis of the up-regulated genes showed that they were mostly mapped to metabolism and virulence pathways, most of which are positively regulated by the global regulator cyclic AMP receptor protein (CRP). Similarly, the transcription level of the *crp* gene in the *pnp*-deletion strain increased 3.22-fold in the early stationary phase. We further explored the indicators of cellular metabolism of the *pnp*-deletion strain, the persistence phenotype of the *pnp* and *crp* double-deletion mutant, and the transcriptional activity of *crp* gene. Our results indicate that PNPase controls cellular metabolism by negatively regulating the *crp* operon at the post-transcriptional level by targeting the 5’- Untranslated Region (UTR) of the *crp* transcript. This study offers new insight about the persister mechanisms and provides new targets for development of new drugs against persisters for more effective treatment of persistent bacterial infections.

## INTRODUCTION

Persisters are a small fraction of dormant or non-growing bacteria that are tolerant to lethal antibiotics or stresses (1). Persisters are distinct from antibiotic-resistant cells in that they are genetically identical and remain susceptible to antibiotics when they are growing again (2, 3). Persisters pose significant challenges for the treatment of many chronic and persistent bacterial infections, such as tuberculosis, urinary tract infections and biofilm infections (4–6). Therefore, it is of great importance to understand the mechanisms of persistence and develop new strategies to more effectively cure such persistent infections. Although the phenomenon of bacterial persistence was discovered over 70 years ago (7), our understanding of the genetic basis of persister formation remains incomplete.

Because the mechanisms of persistence are highly redundant, new mechanisms of persister formation and survival are continually discovered. In bacteria, RNA decay, which is necessary for recycling of nucleotides and for rapid changes in the gene expression program is mainly carried out by RNA degradosome (8). PNPase encoded by *pnp* is a major component of the RNA degradosome, which is composed of a complex structure with RNase E, helicase RhlB, and enolase together with PNPase. In bacteria, mRNA degradation is of great significance, as it not only achieves the nucleotide recycling, but also can control gene expression in different growth conditions (9, 10). PNPase has been found to be important in many aspects of RNA metabolism (8, 11–13). In addition, it also plays important roles in post-transcriptional regulation of gene expression (14).

In this study, to address the role of RNA degradation in bacterial persistence, we constructed *pnp* knockout strain and its complementation strain, and then assessed the survival of the *pnp* gene knockout mutant and the complementation strain upon exposure to antibiotics and stress conditions. In addition, RNA-Seq was performed to evaluate the transcriptome of the *pnp* knockout strain compared with the parent strain W3110, and the differential expression was analyzed to shed light on the molecular basis of the *pnp* mediated persistence. We demonstrate that PNPase controls cellular metabolism by negatively regulating the global regulator cyclic AMP receptor protein(CRP) operon at the post-transcriptional level through targeting the 5’-Untranslated Region (UTR) of the *crp* transcript to regulate persister formation.

## MATERIALS AND METHODS

### Bacterial strains and growth media

The strains used in this work were derived from wild type *E. coli* K12 W3110 (F–mcrAmcrBIN(rrnD-rrnE)1 lambda–). Luria-Bertani (LB) broth (0.5% NaCl) and agar were used for bacterial cultivation.

### Construction of *E. coli* W3110 knockout mutants

The λ Red recombination system was used for construction of *pnp* knockout in the *E. coli* chromosome (15). The candidate gene was replaced by the chloramphenicol resistance gene, which can be removed by pCP20. Primers used for knockout and additional external primers used to verify the correct integration of the PCR fragments by homologous recombination are shown in Table 1.

**Table 1.**
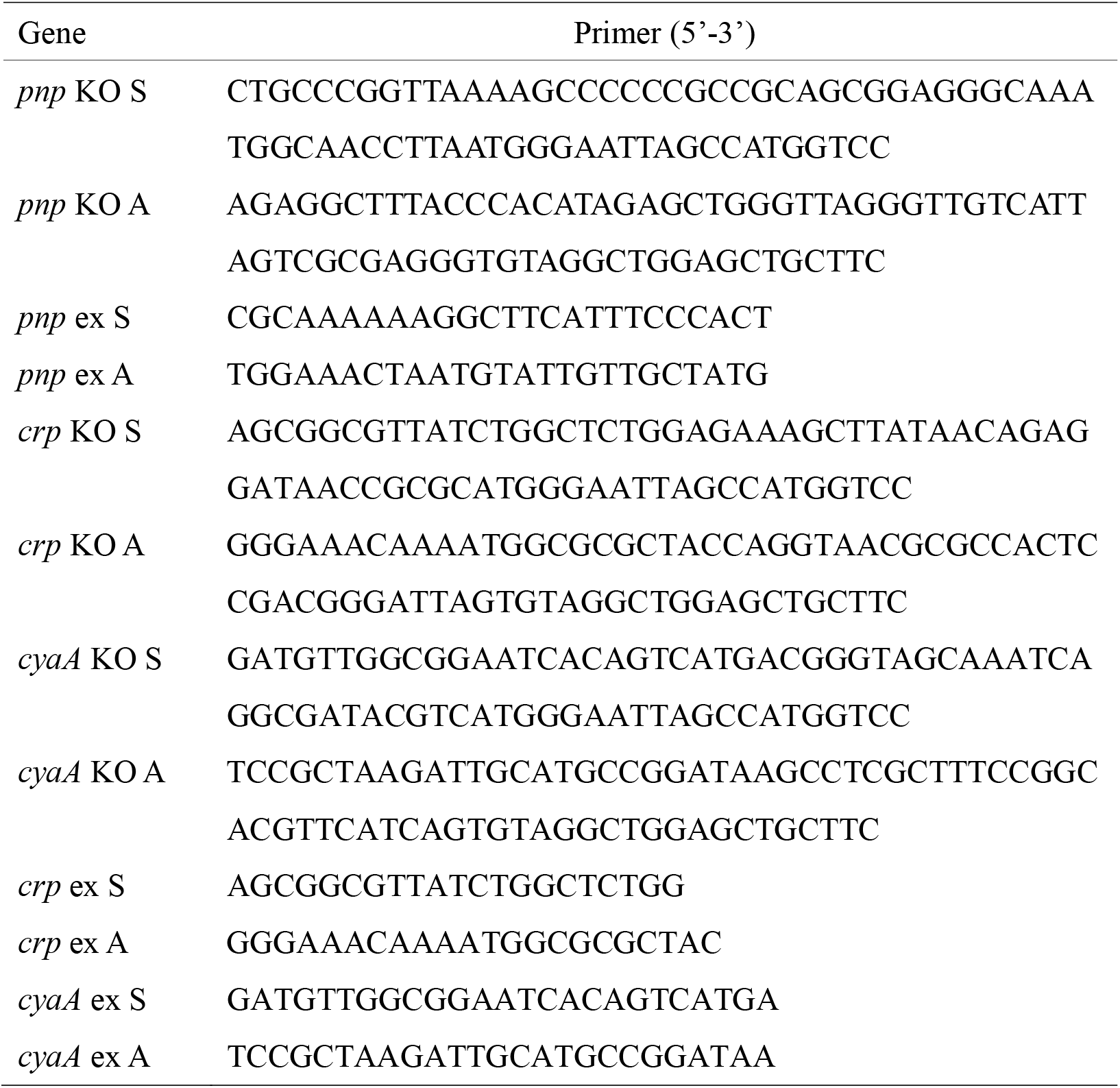
Primers used in construction of *E. coli* W3110 knockout mutants

### Construction of pBAD202-pnp recombinant plasmid

The plasmid pBAD202 was used for construction of a recombinant containing a functional wild type *pnp* gene (16). The primers designed based on the *pnp* gene were F (5’-CATGCCATGGACCCACATAGAGCTGGGTTA-3’) and R (5’-CCCAAGCTTGCAAATGGCAACCTTACT-3’). The PCR products were digested with the restriction enzymes *Nco*I and *Hind*III (Thermo Fisher, USA) and ligated to the plasmid digested with the same enzymes. The recombinant constructs containing the *pnp* gene and the vector control were used to transform the Δ*pnp* deletion strain and wild type W3110 strain by electroporation.

### Persister assay for various antibiotics

Persistence was measured by determining bacterial survival in the form of colony forming units (CFUs) upon exposure to three antibiotics, namely, ampicillin at 200 μg/ml, norfloxacin at 8 μg/ml, and gentamicin at 40 μg/ml. *E. coli* cells were grown to stationary phase in LB medium, and then were exposed to different antibiotics, where undiluted cultures were used for incubation without shaking at 37°C for various times (17). The number of CFUs per milliliter was determined by plating dilutions of the bacterial cells on LB plates without antibiotics.

### Persister assay for various stresses

For heat stress, *E. coli* cells from stationary phase cultures were treated at 52 °C for 30 minutes. The CFUs were determined after serial dilutions. For acid stress, *E. coli* cells were incubated with acid pH 3 for 4 days at 37°C. For hypertonic saline stress, cultures were grown in LB medium containing 3M NaCl at 37°C for 6 days without shaking. *E. coli* cells were also exposed to 80 mM H_2_O_2_ at 37°C for 5 days without shaking. The number of CFUs under acid, hypertonic saline, and H2O2 was determined daily.

### RNA extraction and sequencing

The wild type W3110 cells and the *pnp* mutants were cultivated at 37°C for 6.5 hours to stationary phase. Total RNA was isolated from 1 ml culture, using the RNeasy Mini kit (Qiagen, USA), according to the manufacturer’s instructions. All procedures for RNA sequencing and alignment of the transcriptome were conducted by Oebiotech (Shanghai, China). RNA sequencing was performed using Illumina HiSeq™2000. Raw reads were filtered to remove low quality sequences, adapter sequences, and reads with poly N. The clean reads were subjected to BLAST search by tophat/bowtie2. Differential expression analysis of two samples was performed using the software DEseq. David Bioinformatics Resource 6.7 was used to perform gene ontology and KEGG pathway analysis.

### Detection of bacterial internal redox status

The concentration of intracellular ATP was detected by BacTiter-Glo Microbial Cell Viability Assay Kit (Promega, USA). The intracellular NADH/NAD^+^ ratio was measured as described (18). Carbonyl cyanide m-chlorophenylhydrazone (CCCP), an oxidative phosphorylation inhibitor, was purchased from Sigma-Aldrich (St Louis, MO, USA). CCCP (100 μM) and antibiotics were added to LB as ATP inhibitors in stationary phase. The number of CFUs per milliliter was determined by plating serial dilutions of bacteria on LB plates without antibiotics after 24h incubation at 37°C with 210 rpm shaking.

### Detection of transcriptional activity of cyclic adenosine monophosphate (cAMP) receptor protein (CRP)

The β-galactosidase gene *lacZ* was inserted into the polycloning site of pET-28 vector, and the original T7 promoter was replaced with the promoter region of the *crp* gene or promoter region of the *crp* gene plus 5’-UTR region. The recombinant plasmid constructs pET-lacZ-Pcrp/pET-lacZ-Pcrp+5U were transformed into *E. coli* W3110 and the *pnp* mutant (Δ*pnp* W3110) by electroporation. The activity of β-galactosidase was measured by the β-Gal assay kit (Invitrogen, USA) according to the manufacturer’s instructions. Enhanced BCA protein assay kit (Beyotime Biotechnology, China) was used to determine the concentration of proteins.

## RESULTS

### *pnp* mutant has defect in persistence to various antibiotics

The stationary phase cultures of the *pnp* mutants and the wild-type strain W3110 as a control were exposed to various antibiotics, including ampicillin (200 μg/ml), norfloxacin (8 μg/ml), and gentamicin (40 μg/ml). The results showed that the Δ*pnp* mutant was more susceptible than the parent strain W3110 to all the three antibiotics. Especially upon treatment with gentamicin, the persister levels of the Δ*pnp* mutant decreased significantly after 24h exposure. The effect of ampicillin and norfloxacin on sterilization was similar. The Δ*pnp* mutant was killed by ampicillin and norfloxacin significantly from the second day. Complementation of the Δ*pnp* mutant with the functional *pnp* gene conferred increased persistence to the three antibiotics to the wild-type levels. However, there was no significant change in the persister level between the *pnp* gene overexpression strain and wild-type strain W3110 (see Figure 1).

**Figure1.**
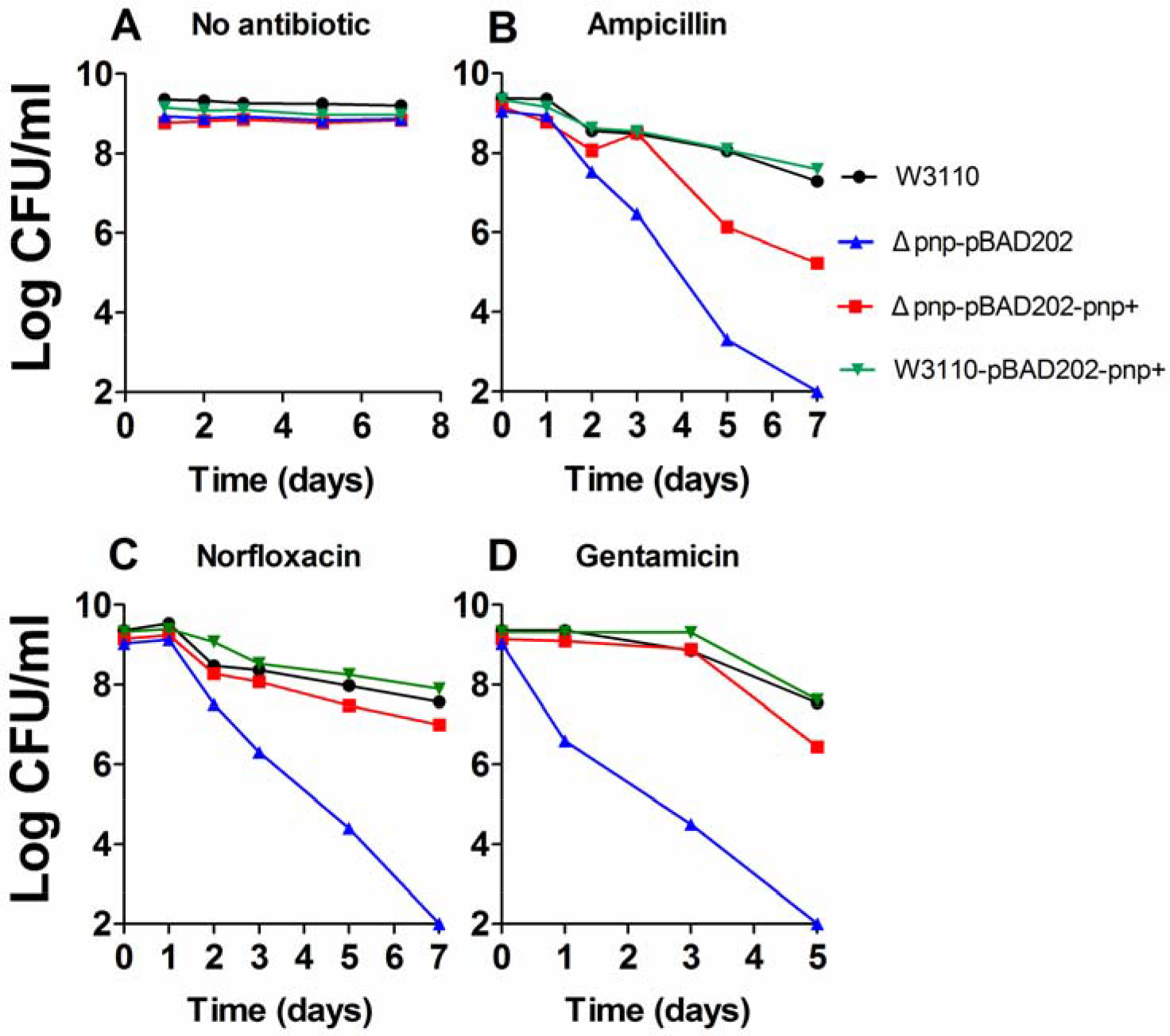
PNPase mutant has defective persister levels in antibiotic exposure assays. Stationary phase culture of the *Δpnp* mutant, its complemented strain, the parent strain W3110, and overexpression strain were exposed to no antibiotic (A), ampicillin (200 μg/ml) (B), norfloxacin (8 μg/ml) (C), gentamicin (40 μg/ml) (D), for various times. Aliquots of cultures were taken at different timepoints and plated for CFU determination on LB plates. The vertical axis represents CFU values on a log scale and the horizontal axis represents time of antibiotic exposure in days.

### The *pnp* mutant is more susceptible to various stresses

Since the persister bacteria are not only tolerant to antibiotics, they can be tolerant to certain stress conditions (3, 19). Thus, we also tested the survival of the Δ*pnp* mutant under several stress conditions, including heat, acid pH, hydrogen peroxide, and hypertonic saline stress. As shown in Figure 2, the stationary phase culture of the Δ*pnp* mutant was more sensitive to the four stress conditions, than that of wild-type strain W3110. Complementation of the Δ*pnp* mutant with the functional *pnp* gene mostly restored the level of persistence of the wild-type strain. It is worth noting that the persistence level was significantly higher in the *pnp* overexpression strains than the wild-type strains under heat stress, but significantly lower in the *pnp* overexpression strains than the wild-type strains under hypertonic saline stress.

**Figure 2.**
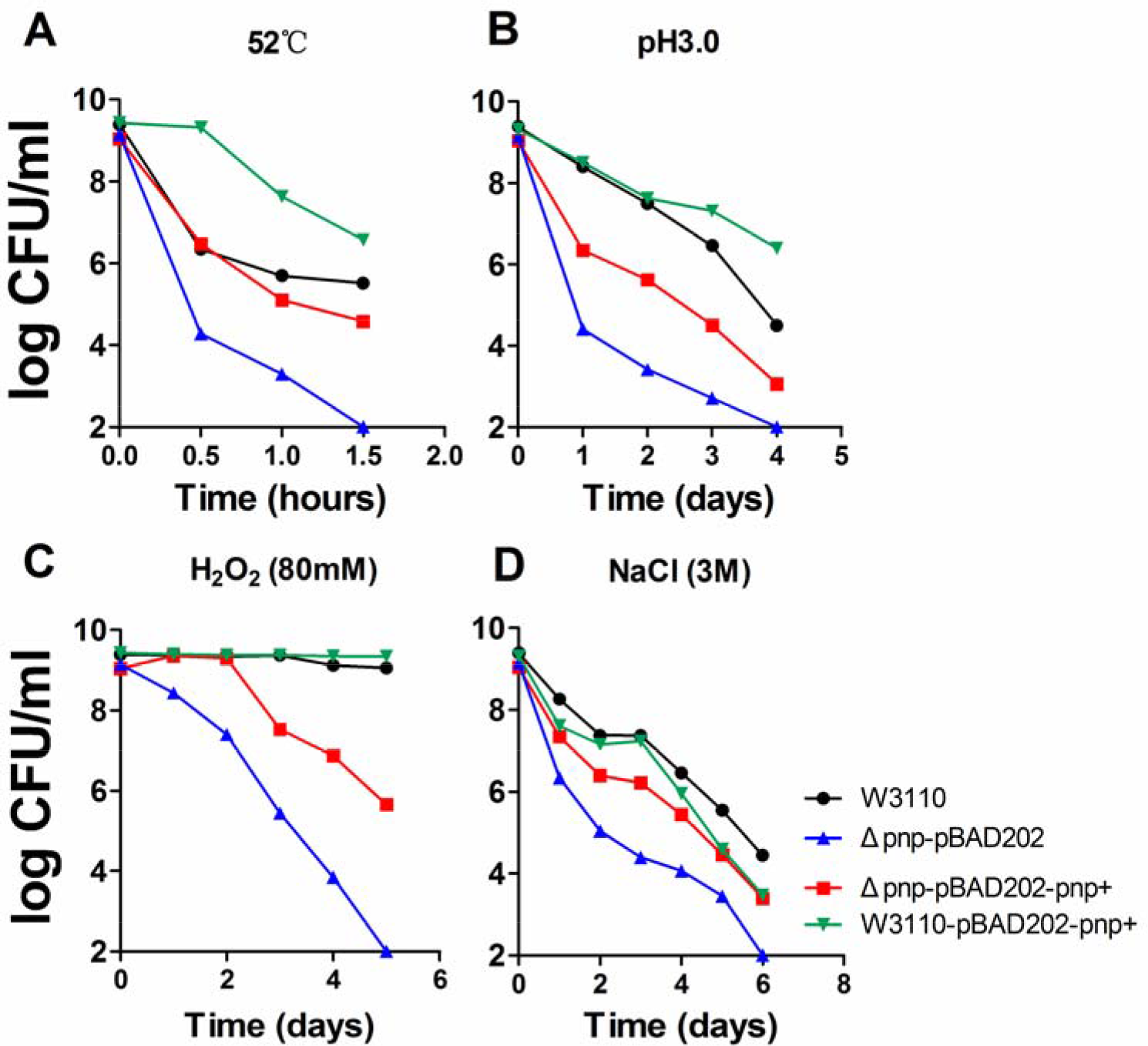
Comparison of susceptibility of four mutants to stresses. The *Δpnp* mutant and its complemented strain, the wild type control W3110, and the overexpression strain to a variety of stress conditions: heat at 52 °C (A); acid pH 3.0 (**B**); hydrogen peroxide (80 mM) (C); NaCl 3M (D). All strains were cultured to stationary phase and exposed to different stresses followed by CFU determination. The vertical axis represents CFU values on a log scale and the horizontal axis represents time of stress exposure in hours or days.

### RNA-seq analysis reveals a higher metabolic status of the *pnp* deletion mutant strain

Figure 3 shows the difference in gene expression profiling between the W3110 wild type strain and the Δ*pnp* mutant. Altogether, 242 genes showed significant differences in the Δ*pnp* mutant compared with the parent strain W3110, where 166 genes were up-regulated and 76 genes down-regulated.

**Figure 3.**
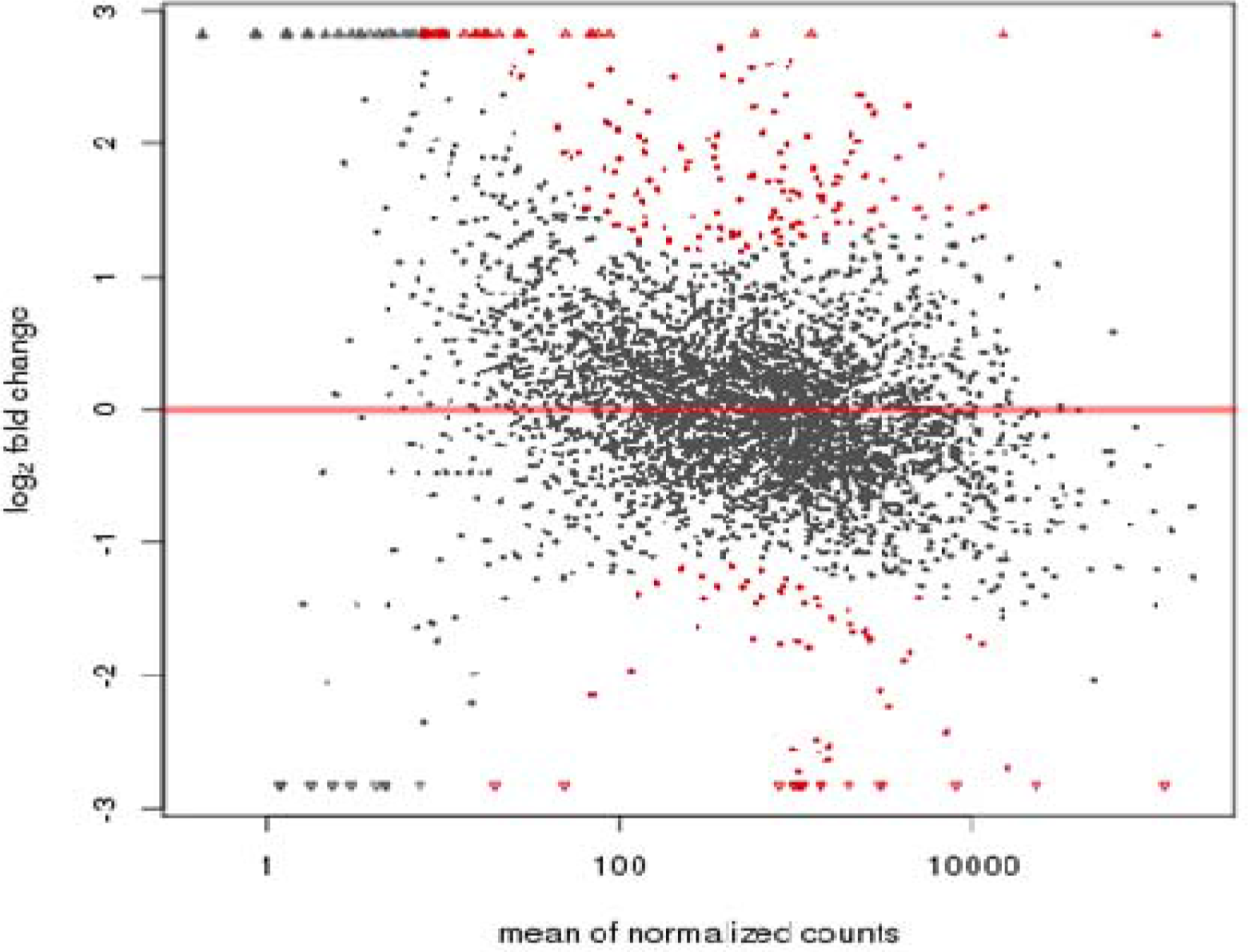
MA map of differential expression of W3110 and the *Δpnp* mutant. Vertical axis represents fold changes in log scale, and horizontal axis represents the mean value of all sample expression for comparison after normalization. Red dots represent the differential genes of p-value<0.05. The transcripts above baseline were up-regulated, and below the baseline were down-regulated.

Pathway analysis of the differentially expressed genes was performed using the KEGG database, and the significance of the differentially expressed gene enrichment in each pathway was calculated by the hypergeometric distribution test. The top 20 pathways after KEGG analysis are shown in Figure 4. Flagellar assembly (PATH: ecj02040), Ribosome (PATH: ecj03010), ABC transporters (PATH: 02010), Sulfur metabolism (PATH: ecj00920), Two-component system (PATH: ecj02020), and carbon metabolism (PATH: ecj01200) are the main enrichment pathways of significant differences in gene transcripts, in which only the genes in ribosome pathway are down-regulated. We selected 41 genes whose mRNA levels increased more than 3 fold in the *pnp* mutant for validation by Real-time quantitative PCR. The results showed that the expression levels of 32 genes increased more than 4-fold (Table 2).

**Figure 4.**
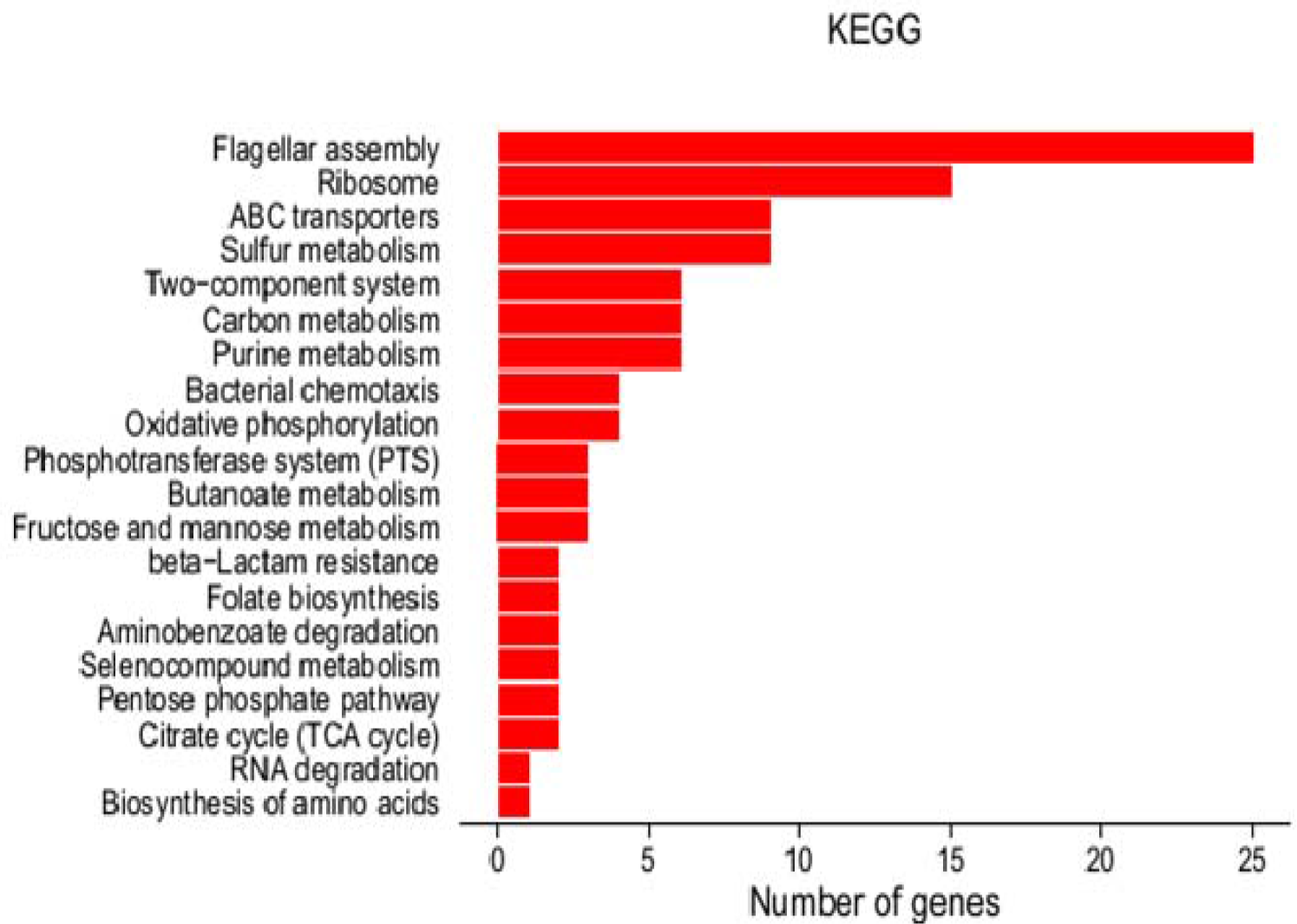
KEGG of differential expression of W3110 and *Δpnp* W3110.

**Table 2.**
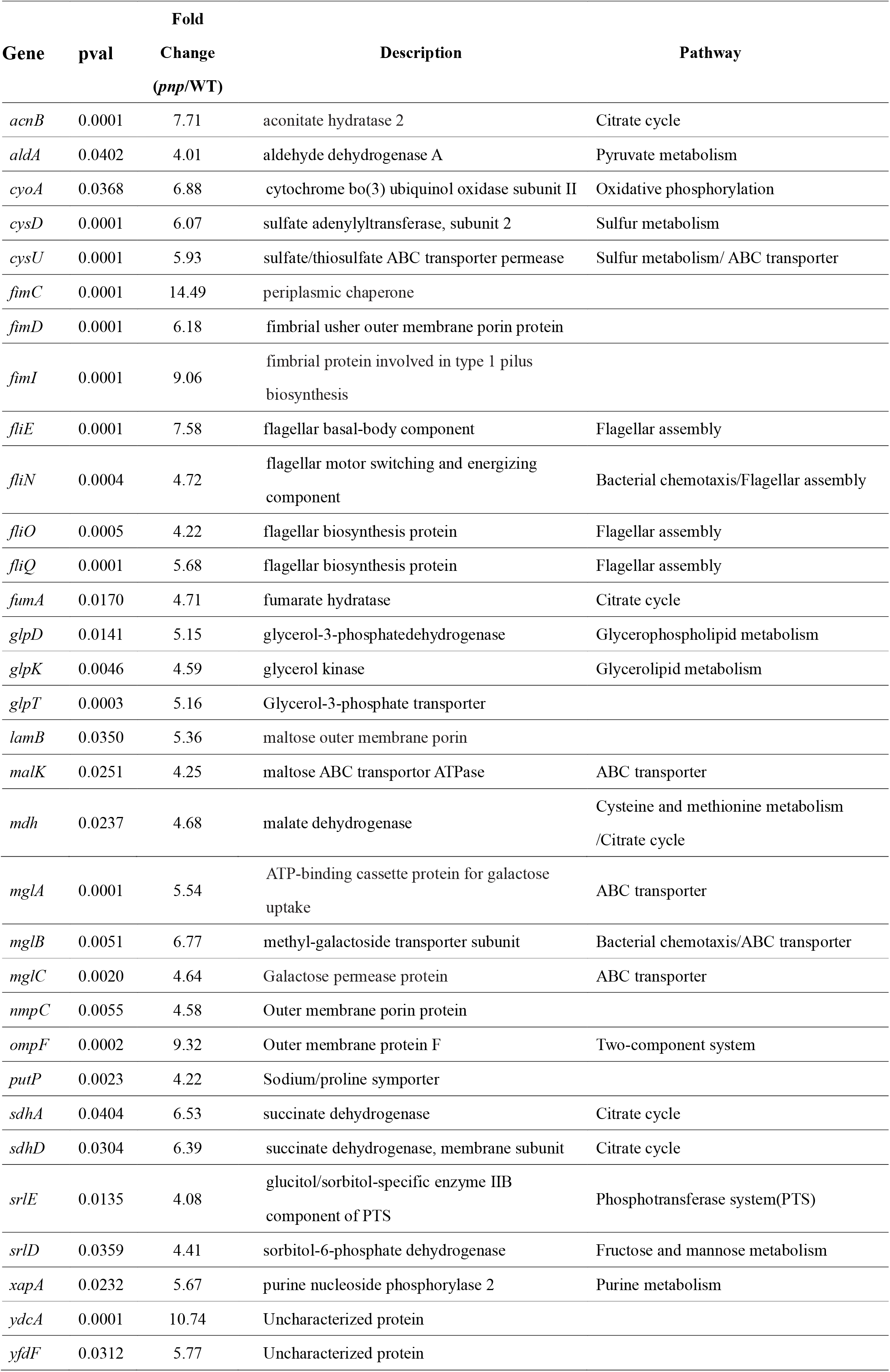
32 genes whose expression ratios were more than 4-fold in *pnp* mutant verified by qPCR

### Detection of internal redox status of bacteria

Since the RNA-seq results showed that genes involved in metabolic and virulence-related pathways were expressed at a higher level in the Δ*pnp* mutant than the parent strain, we measured the intracellular ATP levels and NADH/NAD^+^ ratios to estimate whether the Δ*pnp* mutant was in a state of higher metabolism than the parent strain W3110. Since the persister assay was performed with stationary phase bacteria, we determined the ATP level in Δ*pnp* mutant and the wild-type strain during the stationary phase. As shown in Figure 5, the ATP level of Δ*pnp* mutant was higher than that of the wild type strain W3110 in stationary phase, suggesting that the higher metabolic status in the Δ*pnp* mutant produces excessive ATP and renders the Δ*pnp* mutant less able to form persisters and thus become more susceptible to antibiotics and stresses. Consistent with the above observation, the NADH/NAD^+^ ratio which reflects the redox status of the microbial cells in the Δ*pnp* mutant was higher than that in the control parent strain in stationary phase, suggesting that the Δ*pnp* mutant was indeed in a high metabolic state (Figure 5).

**Figure 5.**
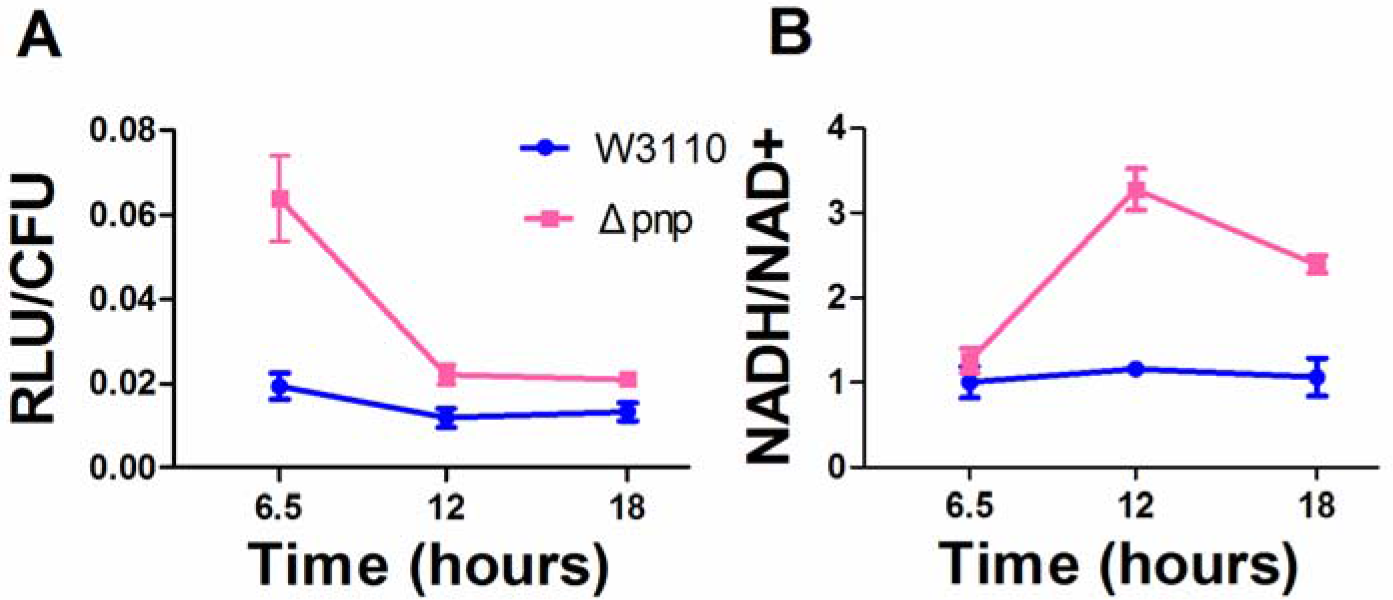
Detection of ATP and NADH/NAD+ in W3110 and the *Δpnp* mutant. (A) The ATP at each stage of the W3110 and the *Δpnp* mutant was detected and the absolute concentration of ATP is calculated by the ATP standard curve. The time points 6.5 h, 12 h and 18 h represent the early stage, the middle stage of stability and the end of stability, respectively. The ATP concentration of the *Δpnp* mutant at all stages is higher than that of the wild type W3110. (**B**) NADH/NAD+ at different stages of W3110 and the *Δpnp* mutant were measured. The *Δpnp* mutant had higher NADH/NAD+ levels than the wild type W3110.

### Antibiotic exposure assays of metabolism related gene knockout strains

Since the RNA-seq analysis indicated that the impact on bacterial metabolism after *pnp* gene deletion is extensive, this suggests that maintenance of the normal metabolism of bacteria by PNPase may be mediated via global regulators. To address this possiblity, we performed real-time quantitative PCR analysis of several major regulators (ArcA, ArcB, Cra, Crp, CyaA, Fnr, and RpoS) responsible for global regulation of diverse aspects of metabolism in *E. coli*. Significantly, we found that in the early stationary phase, expression ofthe *cyaA* encoding adenylate cyclase and *crp* encoding cAMP receptor protein (CRP) increased 3.13 and 3.22 fold, respectively. therefore, we knocked out the *cyaA* and *crp* and assessed their effect on persister levels in drug exposure experiments. In the killing curve experiments (Figure 6), persister levels of the Δ*crp* deletion strain and Δ*cyaA* deletion strain were significantly higher than the wild-type W3110 under gentamicin treatment. The persister levels of the Δ*pnp*Δ*crp* double deletion strain and Δ*pnp*Δ*cyaA* deletion strain under gentamicin and norfloxacin treatment were much higher than that of the Δ*pnp* mutant.

**Figure 6.**
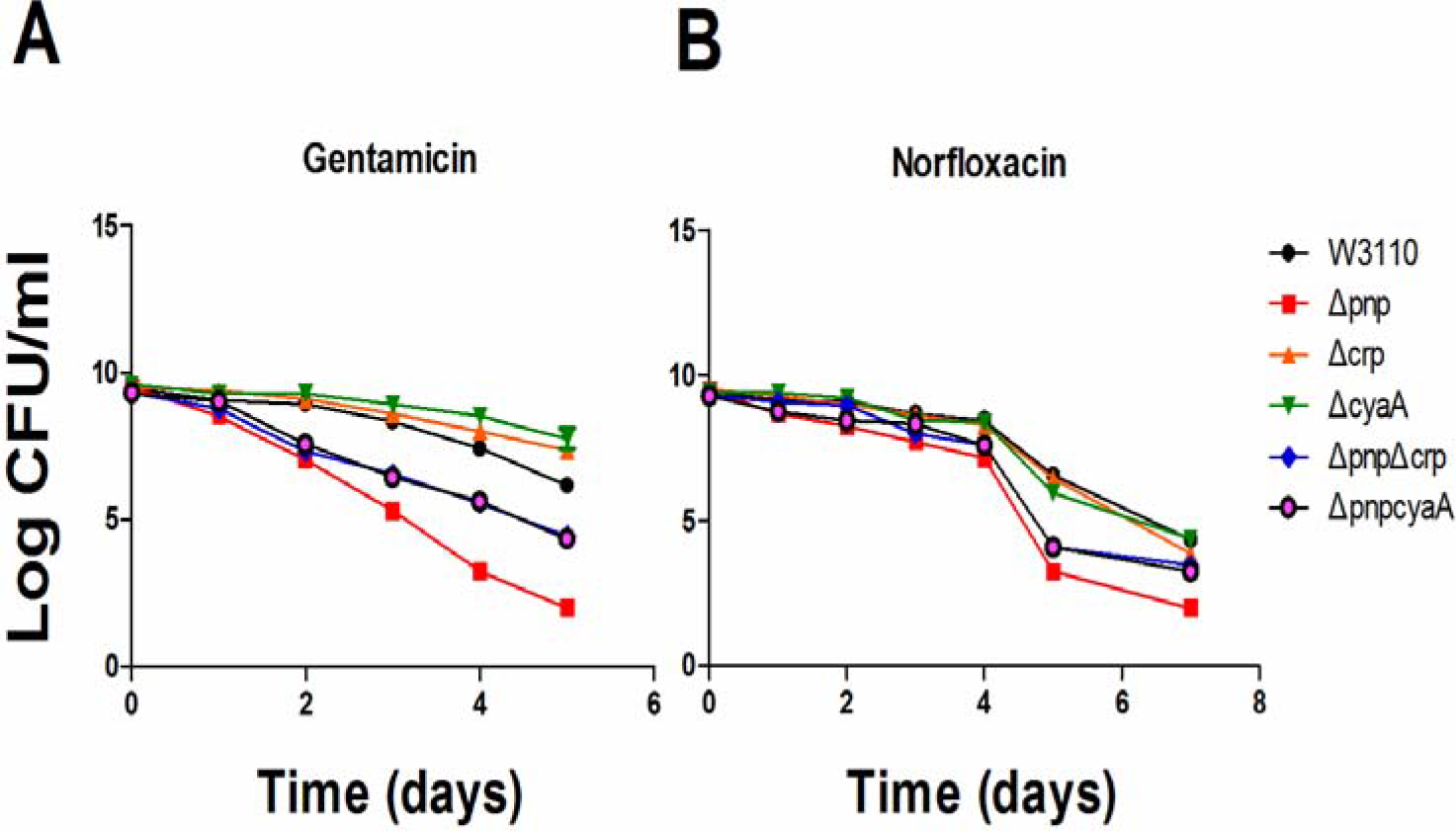
Double KO mutants, *ΔpnpΔcrp* and *ΔpnpΔcyaA* show severe defect in persistence in antibiotic exposure assays. Stationary phase cultures of the control strain W3110 and the mutants *Δpnp, Δcrp, ΔcyaA, ΔpnpΔcrp* and *ΔpnpΔcyaA* were exposed to antibiotics, (A), gentamicin (40 μg/ml) and (B), norfloxacin (40 μg/ml), for various times. Aliquots of cultures were taken at different time points and plated for CFU determination on LB plates. The vertical axis represents CFU values on a log scale and the horizontal axis represents time of antibiotic exposure in days.

### Effect of energy inhibitor CCCP on persister level

In order to further verify the increased sensitivity to antibiotics was caused by high level of metabolism in the bacteria, we performed ATP inhibition experiments. The results showed that addition of CCCP (100 μM) caused significantly higher tolerance to antibiotics. The results are shown in Figure 7.

**Figure 7.**
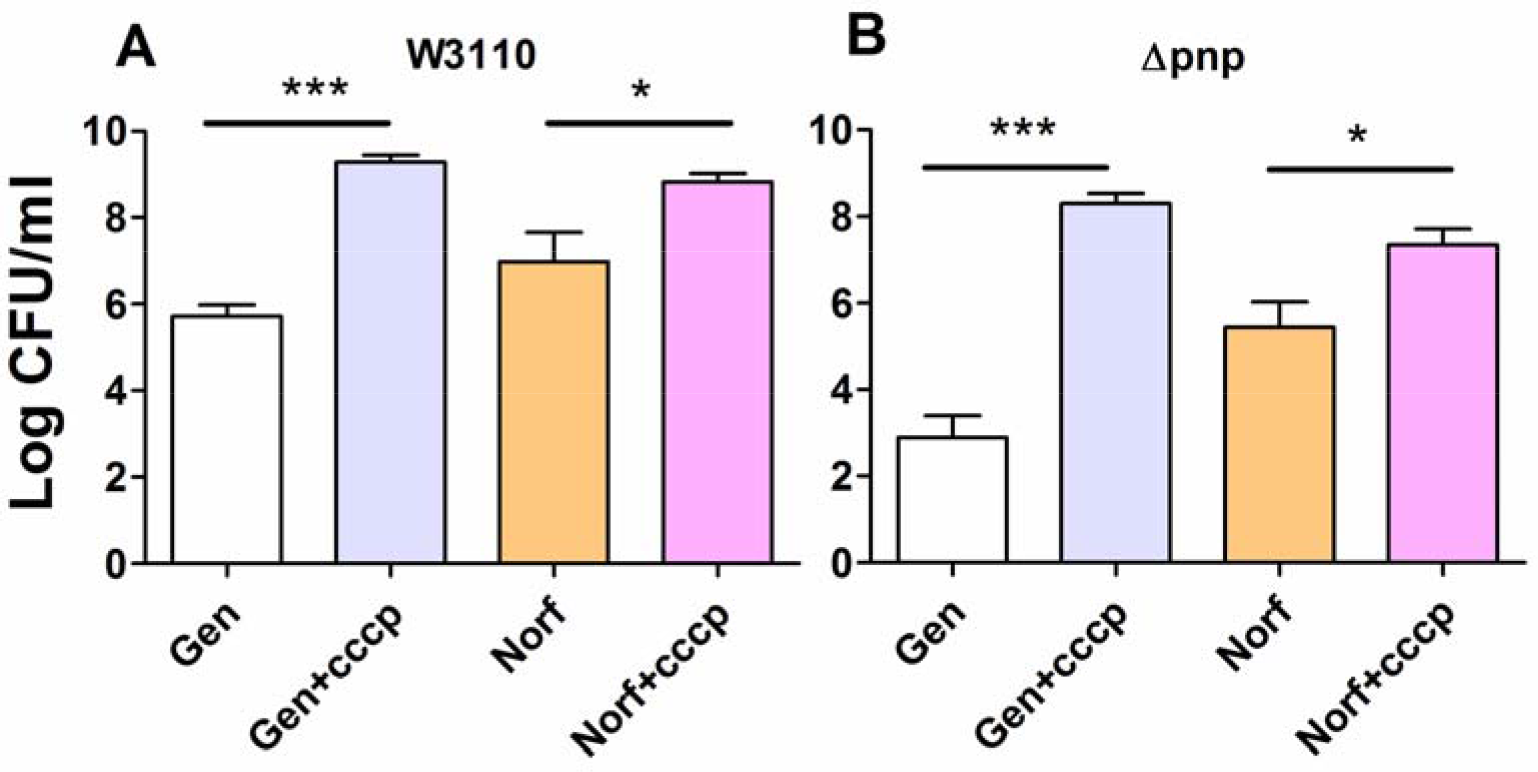
CCCP addition increases the tolerance of W3110 and *Δpnp* mutant to antibiotics. W3110 and the *Δpnp* mutant were exposed to gentamicin (A) or norfloxacin (B) with or without CCCP. CCCP addition significantly increased the persister level. Gen, Gentamicin; Norf, Norfloxacin; CCCP, Carbonyl cyanide m-chlorophenylhydrazone. *t* tests were performed to compare the effect of CCCP addition. *: *P* < 0.05, ***: *P* < 0.0001.

### Transcriptional activity of *crp* gene in W3110 and *pnp* deletion strains

Previous results suggested that PNPase is likely to maintain the downstream metabolic enzymes at the normal level by inhibiting CRP. The 5’-UTR region of prokaryotes is one of the important components of post-transcriptional regulation and can affect the initiation of mRNA translation. It has been found that C-1a PNPase in *E. coli* can bind to 5’-UTR region of the operon *pga*ABCD and inhibit the formation of biofilm by inhibiting the expression of acetylglucosamine (14). Therefore, we hypothesized that PNPase may also bind to the 5’-UTR region of *crp* mRNA to inhibit CRP protein translation. To address this, we made *crp-lacZ* reporter construct and transformed into the Δ*pnp* mutant and the parent strain W3110. As shown in Figure 8, the results of β-galactosidase activity assay showed that the β-galactosidase activity was 8.3-fold higher in the Δ*pnp*/PET-lacZ-Pcrp+5u than that in the parent strain W3110/PET-lacZ-Pcrp+5u at early stationary phase, and 3.6-fold higher at the end of stationary phase. The activity of β-galactosidase in W3110/PET-lacZ-Pcrp was 5.1-fold that in W3110/PET-lacZ-Pcrp+5u at early stationary phase, and 4.1-fold at the end of stationary phase. At any stage, the activity of β-galactosidase in the Δ*pnp*/PET-lacZ-Pcrp was the highest. These findings suggest that PNPase may inhibit the expression of *crp* mRNA or CRP protein translation, leading to higher metabolic status of the Δ*pnp* mutant.

**Figure 8.**
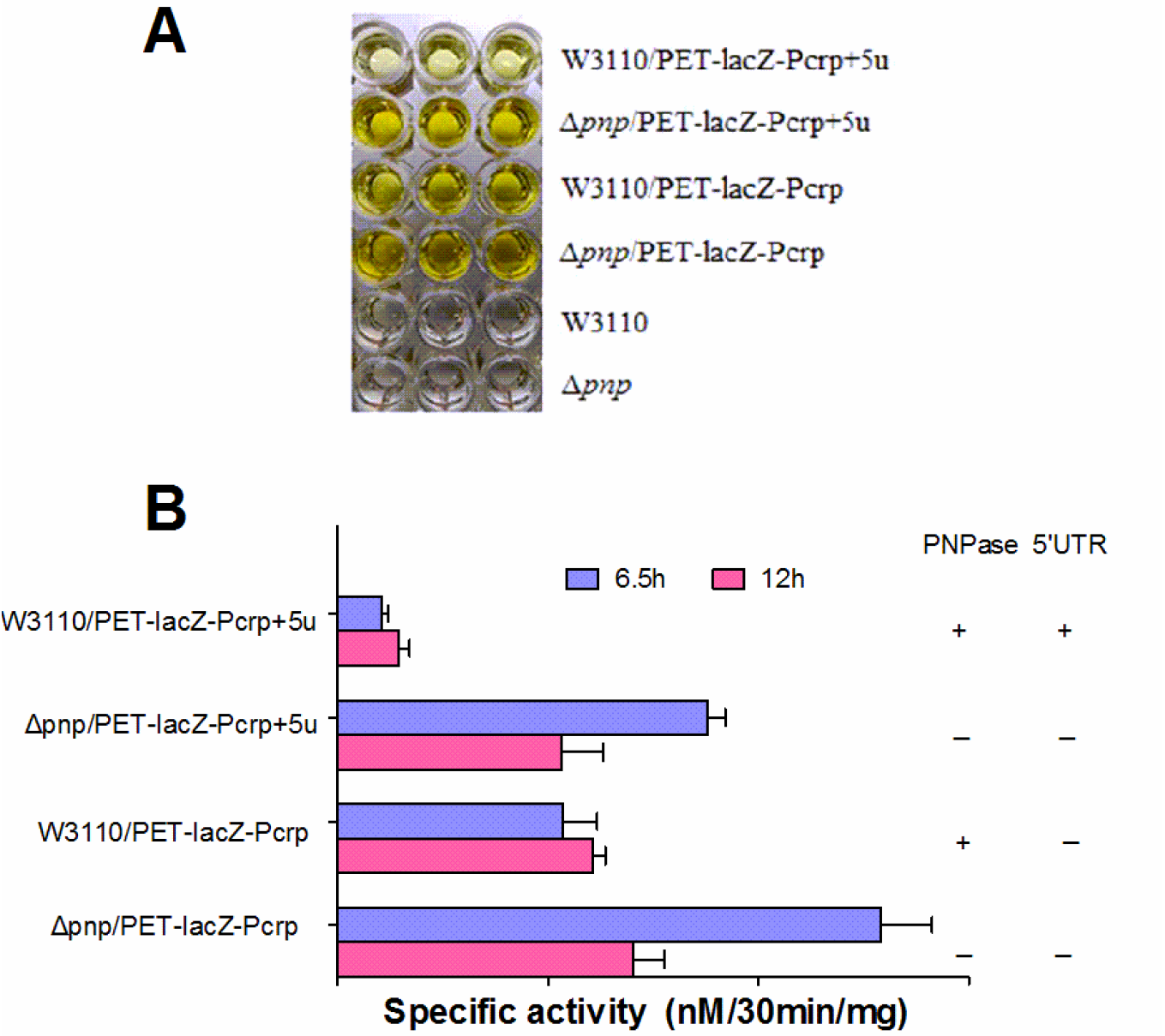
PNPase regulates CRP activity through binding to the 5’-UTR region of *crp* mRNA. (A) The color reaction of β-Gal assay of four strains in 96 well plate. (B) Elevated expression of CRP transcriptional activity (β-Gal specific activity) in the PNPase mutant.

## DISCUSSION

In this study, we identified a new mechanism of persistence mediated by PNPase, part of the RNA degradasome. PNPase catalyzes the polymerization of nucleoside diphosphate and the phosphorylation of polynucleotides in vitro. But in vivo, PNPase is not only the main component of RNA degradasome, which catalyzes decomposition of RNA and promote metabolism and stability of mRNA, but also is independently involved in regulation of bacterial pathogenicity. For example, in *Salmonella enterica*, PNPase acts as a global regulator of virulence gene expression (20).

We found that, compared to wild-type, the Δ*pnp* deletion strain was highly susceptible to three different antibiotics and stresses, indicating that *pnp* plays an important role in the formation or maintenance of persisters. In order to confirm that PNPase was the direct cause of persister formation, we constructed *pnp* complementation strain and found that the persistence phenotype was restored in the *pnp* complementation strain. These *in vitro* experiments have shown that PNPase participates in persister formation or survival. It is worth noting that phenotype of the Δ*pnp* deletion strain mainly appeared in early stationary phase and mid-stationary phase. When the cultures were incubated to late stationary phase, these differences became less obvious, which indicates that the role of PNPase in maintaining the persistence state likely occurs in the early stage.

To shed new insight on the mechanism of by which PNPase mediates persistence, we performed RNA-seq analysis of the Δ*pnp* deletion strain compared with the wild type W3110 strain, and found a number of genes belonging to metabolism or virulence pathway are up-regulated in the Δ*pnp* deletion strain. These finding suggest that elevated metabolism in the Δ*pnp* deletion strain is likely present. This is confirmed by measurement of the ATP levels and NADH/NAD^+^, two indicators of redox state in bacteria, and indicate that the Δ*pnp* deletion strain had significantly higher metabolism than that of the wild type W3110 strain. The *Escherichia coli* CRP is an important transcription factor that its DNA-binding can result in positive or negative regulation of more than 100 genes mainly involved in catabolism of carbon sources other than glucose. In our study, the results of validation of differentially expressed genes showed that the mRNA levels of genes encoding carbohydrate transport systems (*mglBAC, lamB, malK*), TCA cycle (*sdhA, sdhD, fumA, mdh*, and *acnB*), Glycerolipid metabolism (*glpD, glpK, glpT*), etc., were significantly up-regulated. Interestingly, the metabolic master regulator CRP has a positive regulatory effect on these genes (21, 22). In addition, the cAMP-CRP complex is known to be involved in regulation of biofilm formation, quorum sensing systems, and the transcription of the nitrogen regulatory system (23–26). When the culture environment lacks glucose, CRP binds to the effector cAMP, and the activated CRP binds to the TGTGAnnnnnnTCACA sequence near the promoter of the regulatory gene, thereby recruiting RNA polymerase to initiate the transcription of the downstream gene (27). CRP was found to play an important role in the metabolic process in *E. coli*, and the perturbation of cAMP-CRP can significantly diminish the metabolic capabilities of persisters, and can prevent drugs like aminoglycosides from getting into the bacterial cells, thus weakening the ability of such drugs to kill persisters (28).

We knocked out *crp* and *cyaA* gene in Δ*pnp* deletion strain, respectively, and found that the ability of persister formation was significantly higher in the double Δ*crp*Δ*pnp* deletion strain and Δ*cyaA*Δ*pnp* deletion strain than that in the Δ*pnp* deletion strain. PNPase can bind to its own 5’-UTR region to realize the autologous expression regulation by promoting degradation of *pnp* mRNA in the presence of RNase III (29–31). We inferred that PNPase in *E. coli* can inhibit the translation of the *crp* mRNA, which controls the CRP level in the normal range, to maintain the balance of intracellular metabolism. To confirm this inference, we measured the activity of β-galactosidase to reflect the transcriptional activity of *crp* in the wild type W3110 strain and the Δ*pnp* deletion strain. We found that the transcriptional activity of *crp* in the Δ*pnp* deletion strain had 8.3-fold increase in the early stationary phase, indicating that PNPase can inhibit the expression of CRP protein at the transcriptional level.

The metabolic status is the critical basis in the formation of persisters in bacteria (32–34). Although some researchers believe that dormancy is not a necessary condition for the formation of persisters (35), the idea that “persistence can be considered a termination of a metabolic procedure, that is bacteria enters a dormant state” remains the mainstream thinking. Based on our findings, we propose that PNPase in *E. coli* can bind to the 5’-UTR of the *crp* mRNA to cause a negative regulation and maintain a constant level of CRP protein, and thatdeletion of PNPase could cause overexpression of CRP, the downstream regulation of a series of metabolic-related gene expression levels increased abnormally, leading to failure to enter the dormant state, causing decreased persistence capacity in the Δ*pnp* deletion strain and its inability to tolerate antibiotics and stress conditions.

In summary, our study established a new mechanism of persister formation mediated by PNPase regulation in *E. coli* (Figure 9). We found that PNPase controls cellular metabolism by negatively regulating the global regulator cyclic AMP receptor protein (CRP) operon at post-transcriptional level through targeting the 5’-UTR region of the *crp* transcript to regulate persister formation. The results of the *in vitro* studies need to be further tested in animal models to determine if PNPase has any role in persistence and virulence in vivo in the future. PNPase, as a regulator of persistence and virulence, is a promising new drug target for developing improved treatment of persistent bacterial infections.

**Figure 9.**
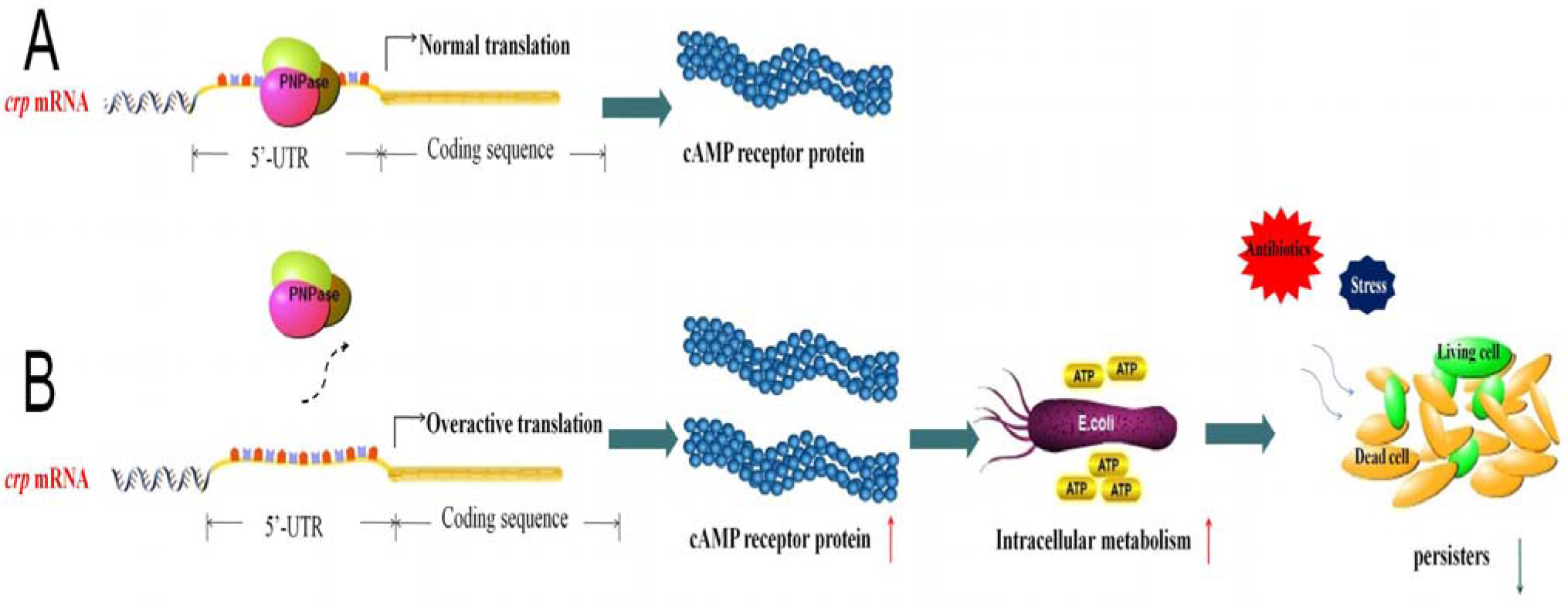
PNPase regulates formation of bacterial persisters through CRP. (A) PNPase binds to the 5’UTR of *crp* mRNA. (B) PNPase deletion leads to higher level of metabolism as seen by elevated redox (NADH/NAD^+^ ratio) and ATP levels and decreased number of persisters.

## ACKNOWLEDGEMENTS

This work was supported in part by the National Natural Science Foundation of China (81572046, 81772231).

